## Supplementary material for "Most primary olfactory neurons have individually neutral effects on behavior": Figures Only

### *Figures and Tables*

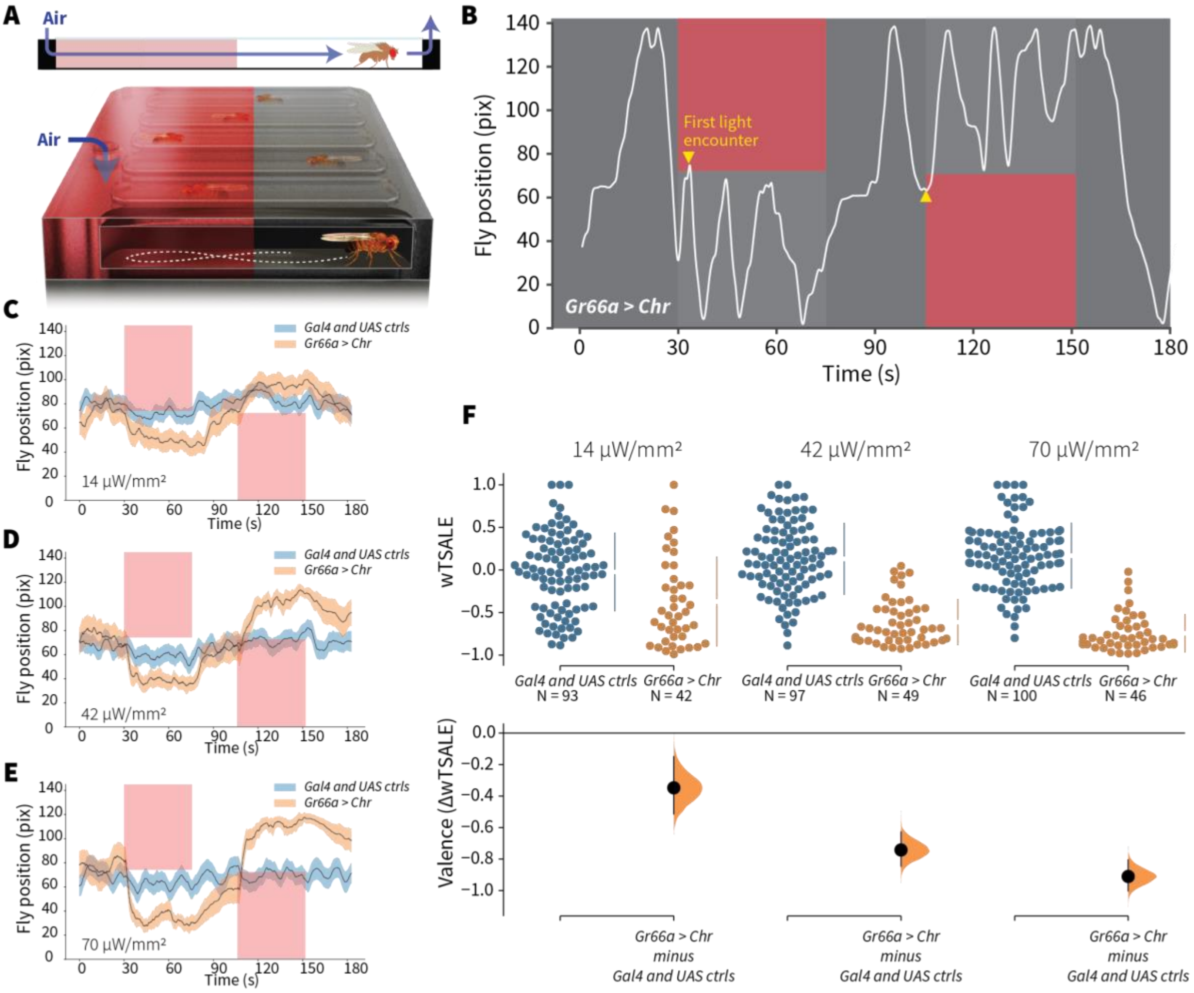


**Figure 1. An optogenetic behavior assay reports on sensory valence**

**A.** Side view schematic (top) showing air flow path and overview (bottom) of the WALISAR assay. Individual flies were placed in chambers and given a choice between no light or red-light illumination. Air flow was used in some experiments.

**B.** A representative path of a fly displaying strong light avoidance. The white line represents the location of a *Gr66a-Gal4 > UAS-CsChrimson* fly in the chamber throughout an experiment. The light preference of each fly was calculated by how much time it spent in the illuminated zones after the initial encounter with light (yellow arrows).

**C-E.** Trace plots representing the mean location of controls ($w^{1118}$; *Gr66a-Gal4* and $w^{1118}$; *UAS-CsChrimson*), and test flies (*Gr66a-Gal4 > UAS-CsChrimson*) throughout an experiment at 14, 42, and 70 µW/mm$^2$ light intensities. The blue and orange ribbons indicate 95% CIs for the control and test flies, respectively.

**F.** An estimation plot presents the individual preference (upper axes) and valence (lower) of flies with activatable *Gr66a+* neurons in the WALISAR assay. In the upper panel, each dot indicates a preference score (wTSALE) for an individual fly: $w^{1118}$; *Gr66a-Gal4* and $w^{1118}$;

*UAS-CsChrimson* flies are colored blue; and *Gr66a-Gal4 > UAS-CsChrimson* flies are in orange. The mean and 95% CIs associated with each group are shown by the adjacent broken line. In the bottom panel, the black dots indicate the mean difference (ΔwTSALE) between the relevant two groups: the valence effect size. The black whiskers span the 95% CIs, and the orange curve represents the distribution of the mean difference.

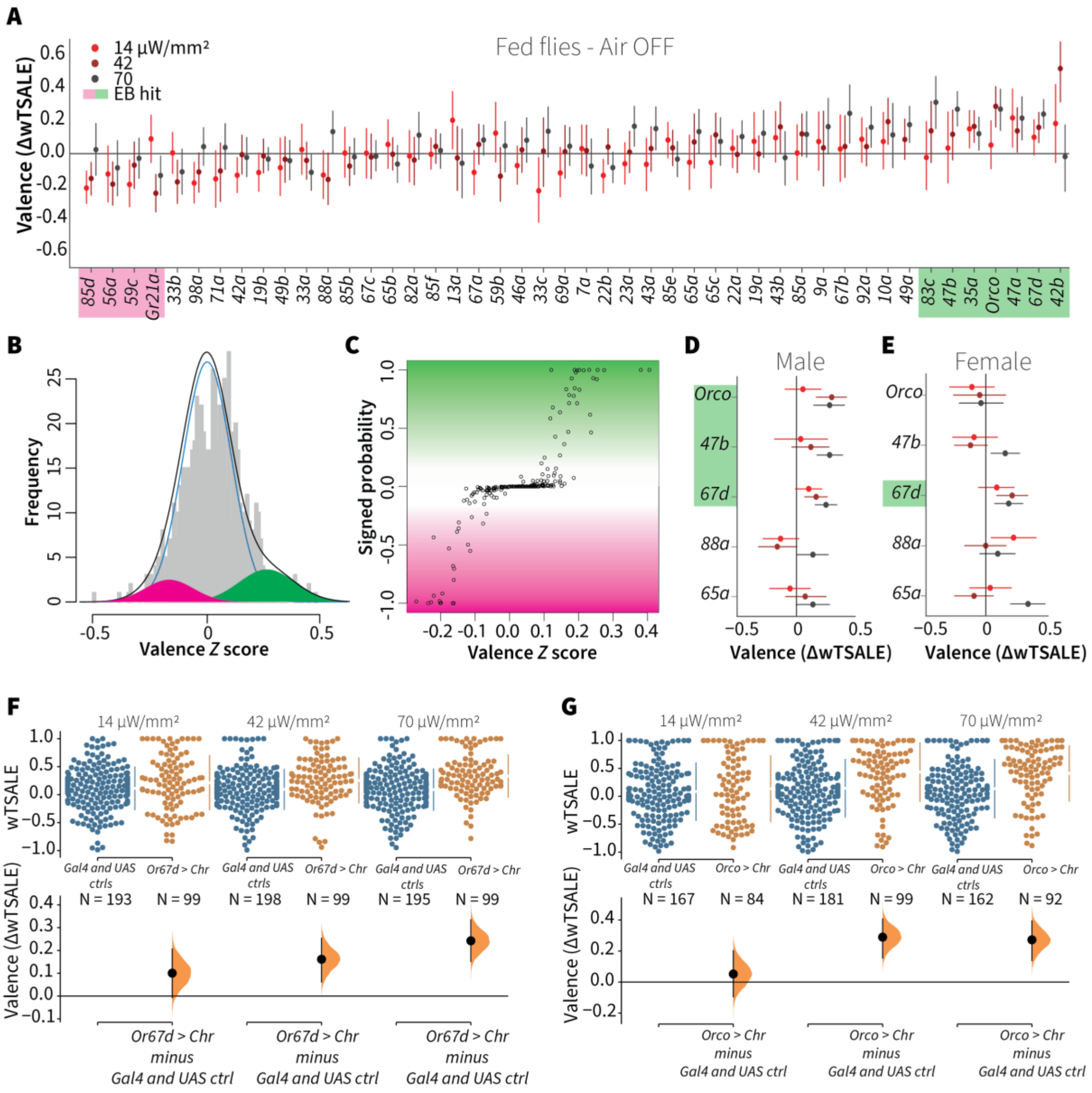


**Figure 2. A small minority of ORN classes individually drive valence**

**A.** An effect-size plot of the valence screen of 45 single-ORN types (and *Orco* neurons). Each dot represents a mean wTSALE difference of control (N ≅ 104) and test (N ≅ 52) flies, whisker indicate 95% CIs. The shades of red represent the three light intensities. Valent ORNs are shaded with magenta (aversion) or green (attraction).

**B.** The histogram (grey bars) of the median ORN ΔwTSALE ratios, and the mixture model that is fit to the data by empirical Bayes method. The blue line represents the null distribution, while the magenta and green curves represent ORNs with negative and positive valences, respectively. The black line represents the overall distribution of the effect sizes.

**C.** The signed posterior probability that each valence score is a true behavioural change, plotted against the respective effect size (median effect across the three light intensities).
**D-E**. The olfactory valence of the *Orco* and four single-ORNs were tested in female flies (the male fly data is replotted from Panel A for comparison). The valence responses are represented as the mean difference (ΔwTSALE) of control (N ≅ 104) and test (N ≅ 52) flies, along with 95% CIs. Color key is the same as above.
**F-G.** Estimation plots show the optogenetic preference (upper panel) and valence (lower panel) of flies with activatable *Or67d* and *Orco* neurons across three light intensities. The dots in the top panels represent single flies, while the broken lines indicate the mean and 95% CIs. The differences between the pairs of test and control groups are displayed in the bottom panel, where the whiskers are 95% CIs and the curves are the distributions of the mean difference.

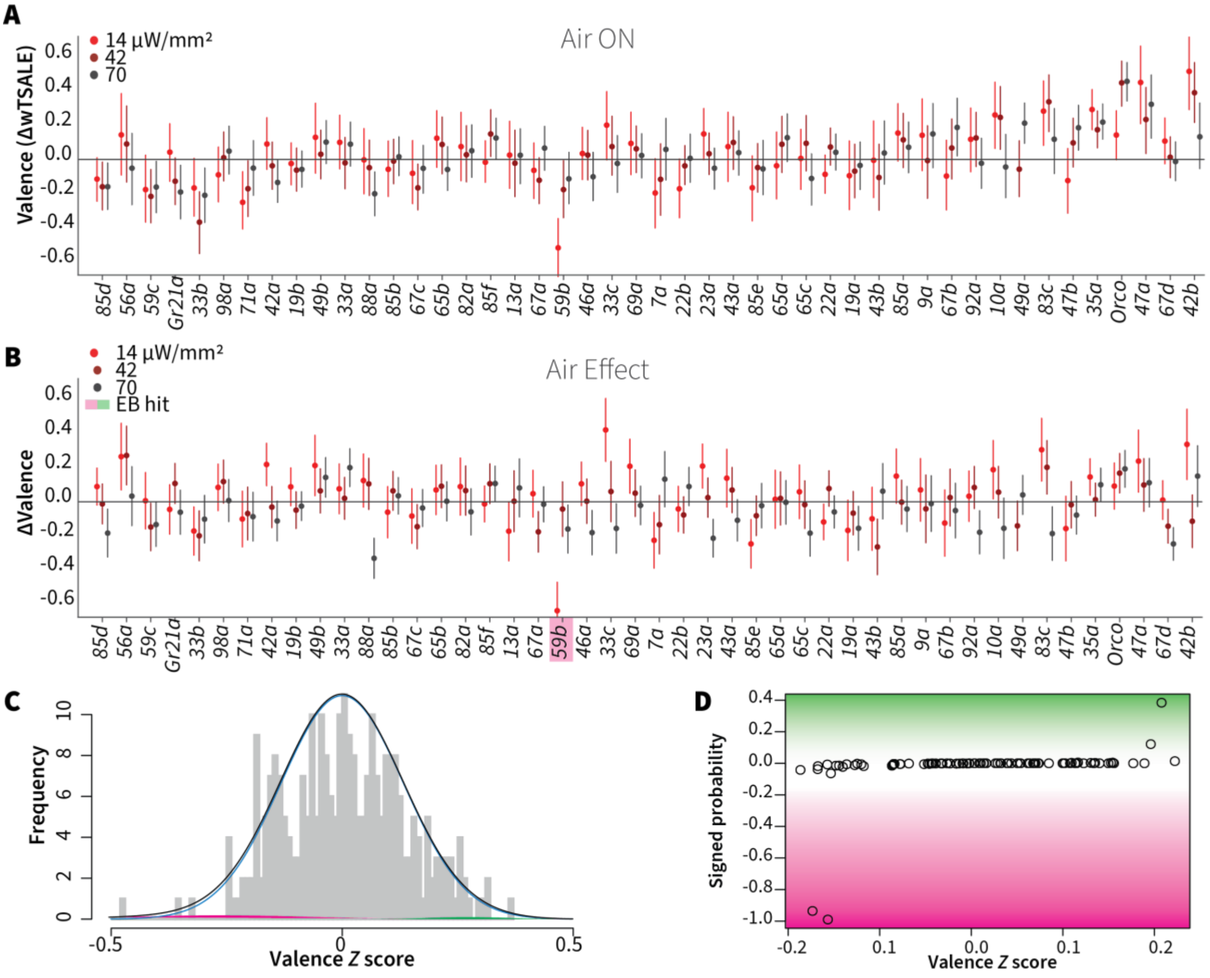


**Figure 3. Wind does not amplify single-ORN valences**

**A.** The results of ORN valence assays in the presence of airflow. The red dots represent the mean wTSALE differences of control (N ≅ 104) and test (N ≅ 52) flies with the 95% CIs.

**B.** The differences between the effect sizes of air-on and air-off experiments (ΔΔwTSALE). The magenta shaded box indicates the sole empirical Bayes hit, Or59b, that showed an increase in aversion at the lowest light intensity only.

**C.** A statistical mixture model was fitted to the ΔΔwTSALE scores. The grey bars are the response histogram. The magenta and green curves represent the effect sizes that differ from the null distribution (blue line). The black line represents the overall distribution of the ΔΔwTSALE scores.

**D.** The signed posterior probabilities of the ΔΔwTSALE scores being true behavioural changes versus their median effect sizes. The overall probability of true wind effects was nearly zero [$p$(ΔΔwTSALE) ≅ 0.0].

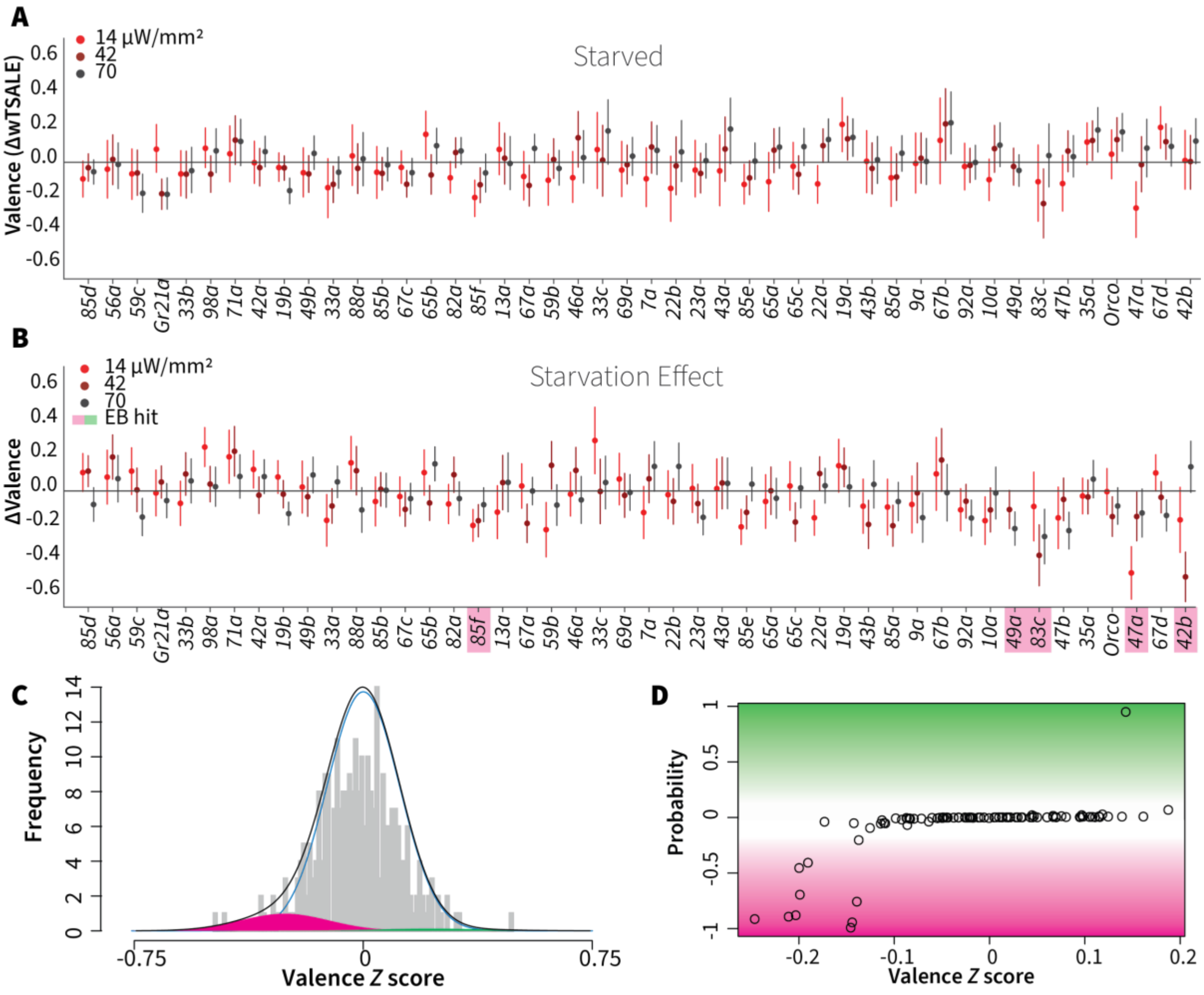


**Figure 4. Starvation affects valence responses for five ORN classes**

**A.** An ORN valence screen for starved animals. The red dots indicate the mean wTSALE differences between control (N ≅ 104) and test (N ≅ 52) flies.

**B.** The mean differences between fed and starved flies across ORNs. The magenta shading indicates the ORNs that are affected by starvation according to the Empirical-Bayes analysis.

**C.** A histogram of the median ΔΔwTSALE ratio distribution. The blue line represents the null distribution, the magenta and green curves represent the distributions of negative and positive valences that are separated from the null distribution, and the black line represents the overall ΔΔwTSALE distribution.

**D.** The signed posterior probability of the ΔΔwTSALE scores being true behavioral changes are plotted against their median ratios.

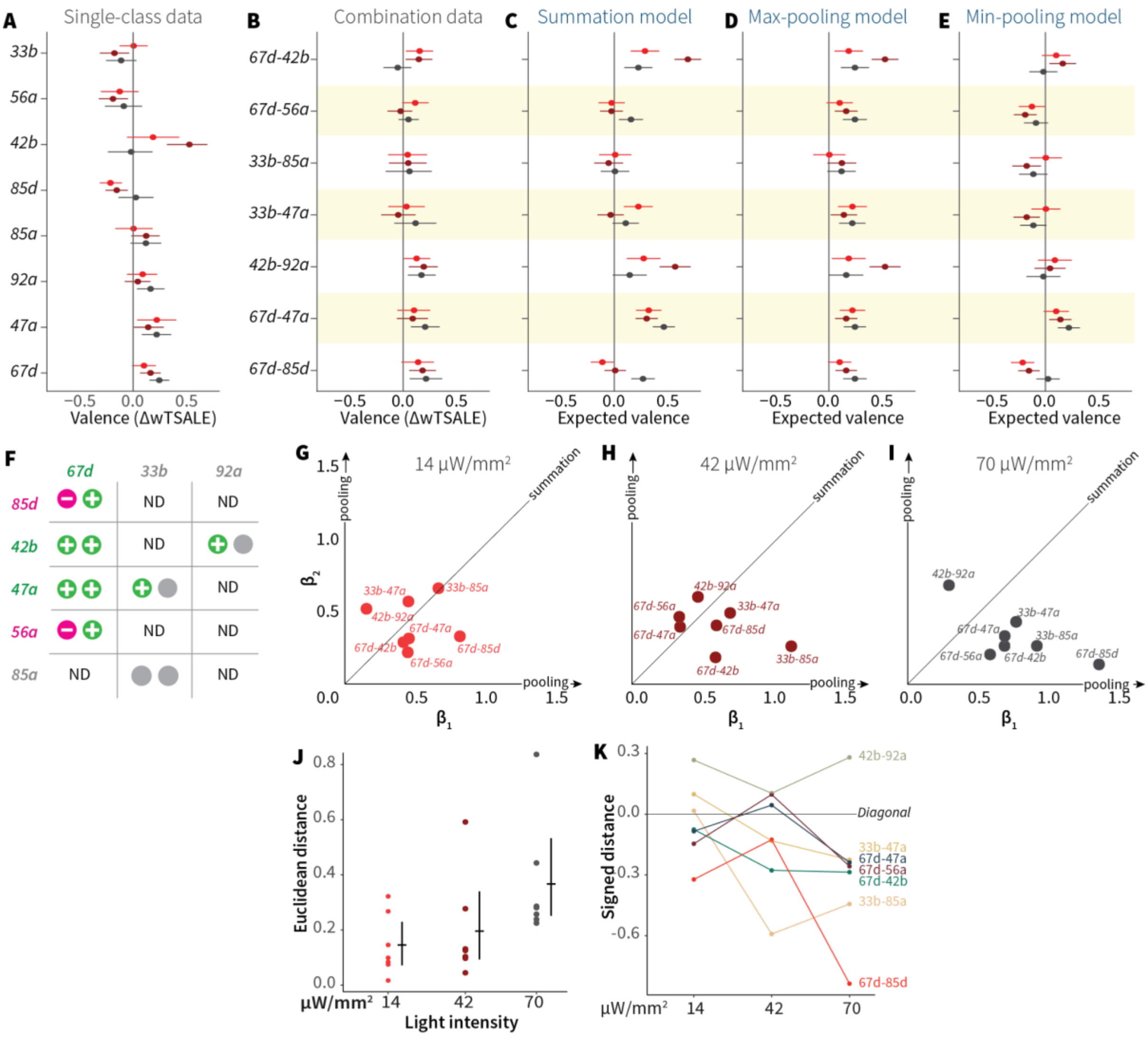


**Figure 5. ORN-valence combinations follow complex rules**

**A.** Valence responses of the single-ORN lines used to generate ORN-combos (replotted from Figure 1). The dots represent the mean valence between control (N ≅ 104) and test (N ≅ 52) flies (ΔwTSALE with 95% CIs). The shades of red signify the three light intensities.

**B.** The valence responses produced by the ORN-combos in the WALISAR assay in three light intensities.

**C-E.** ORN-combo valences as predicted by the summation (C), max pooling (D) and min pooling (E) models.

**F.** Three positive (green), two negative (magenta), and three neutral (grey) ORNs were used to generate seven ORN two-way combinations.

**G-I.** Scatter plots representing the influence of individual ORNs on the respective ORN-combo valence. The red (G), maroon (H), and black (I) dots indicate ORN-combos at 14, 42, and 70 μW/mm² light intensities, respectively. The horizontal ($β_1$) and vertical ($β_2$) axes show the median weights of ORN components in the resulting combination valence.

**J.** Euclidean distances of the ORN-combo β points from the diagonal (summation) line in panel J. The average distance increases as the light stimulus intensifies: 0.14 [95CI 0.06, 0.23], 0.20 [95CI 0.06, 0.34], and 0.37 [95CI 0.20, 0.53], respectively.
**K.** The β weights of the ORN-combos from the multiple linear regression are drawn as the signed distances of each ORN-combo from the diagonal line over three light intensities. The ORN weightings change magnitude and, in a few cases, the dominant partner changes with increasing optogenetic stimulus.

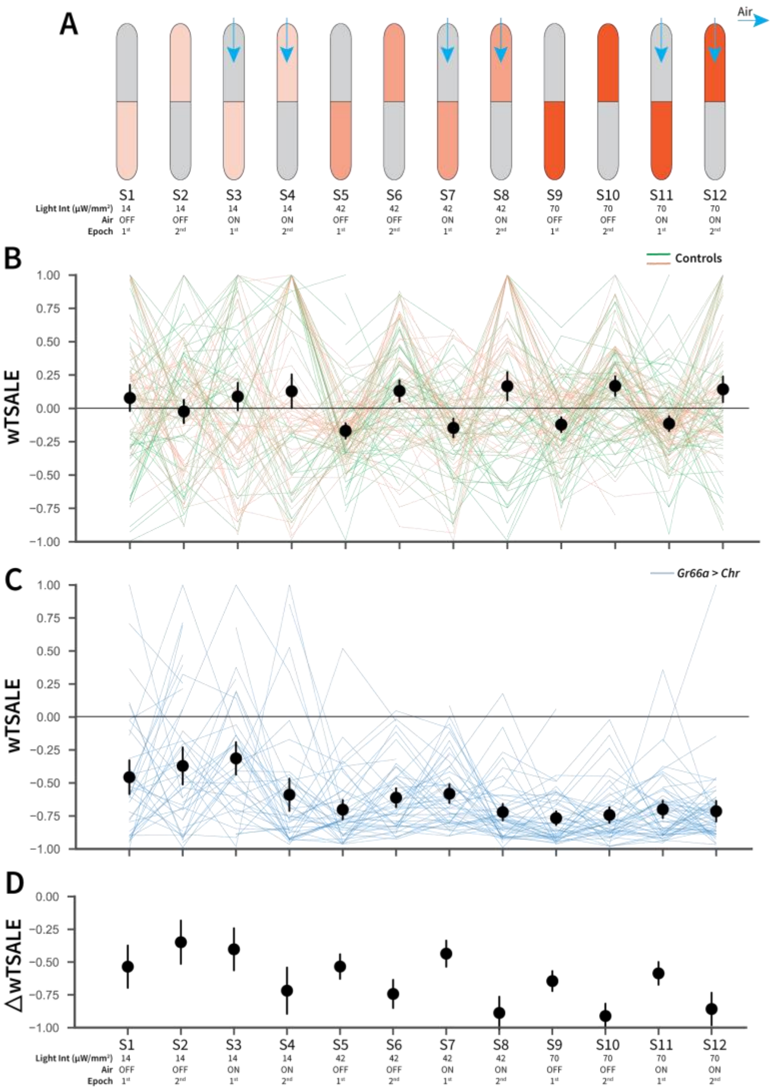


**Figure 1—figure supplement 1. An optogenetic behavior assay reports on Gr66a-induced avoidance**

**A.** In each experiment, the flies were subjected to 12 consecutive epochs: three light intensities that were applied in an increasing order (14, 42, 70 µW/mm$^2$); with and without airflow; and the illumination side was flipped in alternating epochs.

**B.** The light preferences of control flies across the 12 epochs, as estimated by weighted time spent after light encounter (wTSALE). Each line represents a single fly; the green lines indicate *$w^{1118}$; Gr66a-Gal4* flies, and the orange lines indicate *$w^{1118}$; UAS-CsChrimson* flies. The black dots represent the mean wTSALE scores of all control flies per condition; the whiskers indicate the 95% CIs.

**C.** The light preference of *Gr66a-Gal4* > *UAS-CsChrimson* flies across the epochs.

**D.** The optogenetic effect sizes, calculated by taking the difference of test and control preferences (ΔwTSALE).

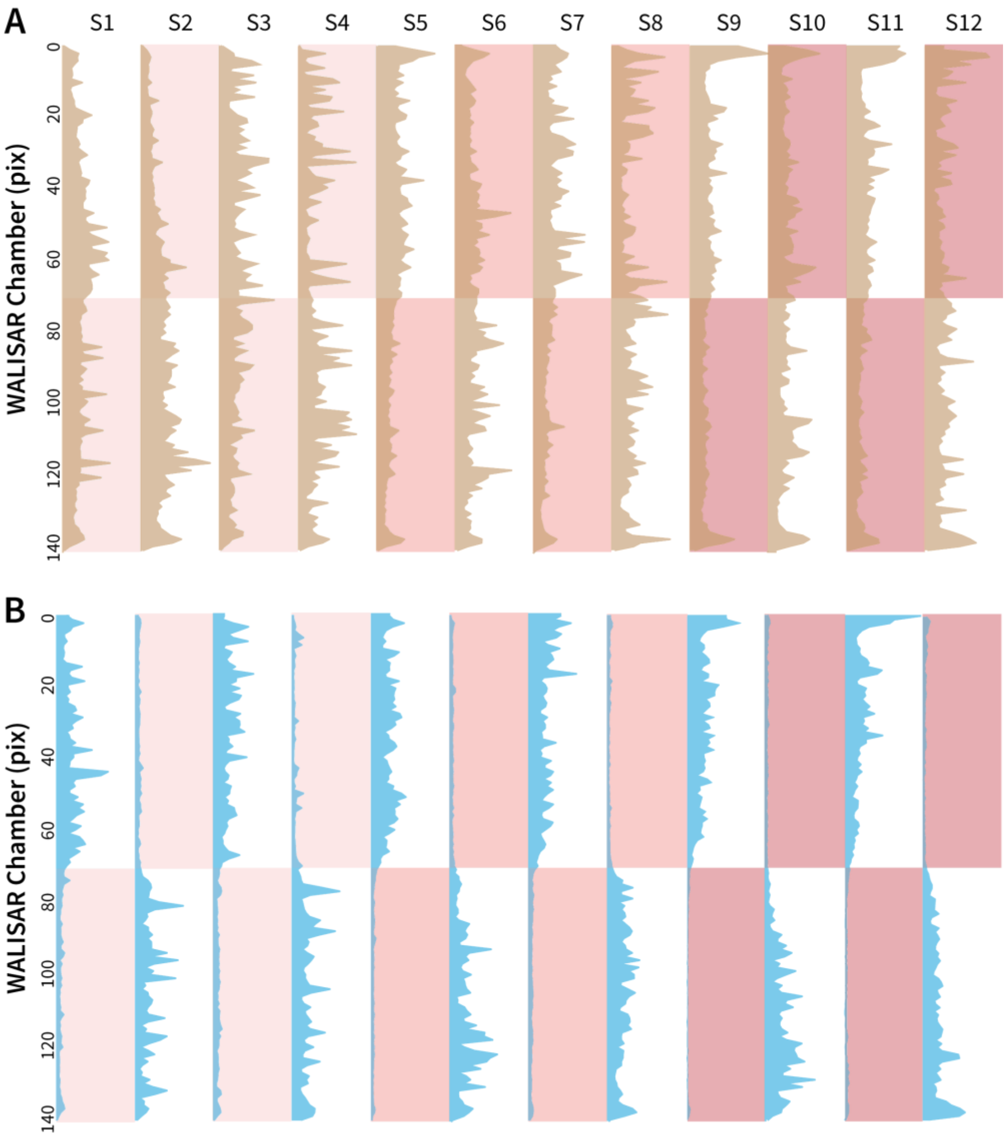


**Figure 1—figure supplement 2. Occupancy analysis of the WALISAR chambers during Gr66a-neuron activation**

**A.** The number of occurrences of the control flies (*w*[1118]*; Gr66a-Gal4* and *w*[1118]*; UAS-CsChrimson*, N ≅ 104) across the WALISAR chambers are drawn. The brown curves present the total number, the y axis represents the chamber, and the x axis is the frequency of occurrence (ranging between 0 and 2,000). The red shading indicates the three light intensities; and the labels from S1 to S12 represent the 12 experimental epochs (Figure 1C). **B.** The number of occurrences of the experimental flies (*Gr66a-Gal4>UAS-CsChrimson*, N ≅ 52) across the WALISAR chambers are summed and drawn for the 12 epochs. The blue curves represent the total number, the y axis represents the chamber, and the x axis is the frequency of occurrence (ranging between 0 and 1,500).

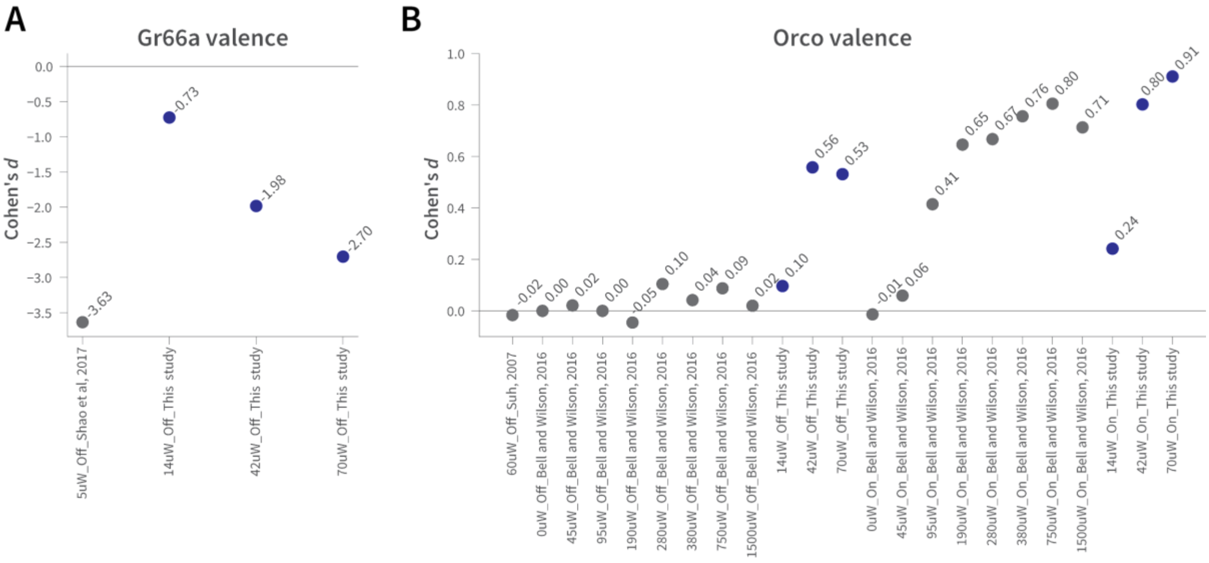


**Figure 1—figure supplement 3. The effect sizes in WALISAR and previously reported optogenetic valence assays are comparable**

**A.** A comparison of the Gr66a-mediated aversion (Cohen's *d*) between this study at different light intensities and Shao et al. at 5 μW/mm$^2$ of blue light, both in still-air conditions (Shao et al., 2017).

**B.** The magnitude of the attraction (standardized effect size, Cohen's *d*) produced by *Orco*-neuron activation in the present study is compared to that of two previous studies (Bell and Wilson, 2016; Suh et al., 2007). The light intensity is shown in μW/mm$^2$, while "On" and "Off" indicate the presence of wind.

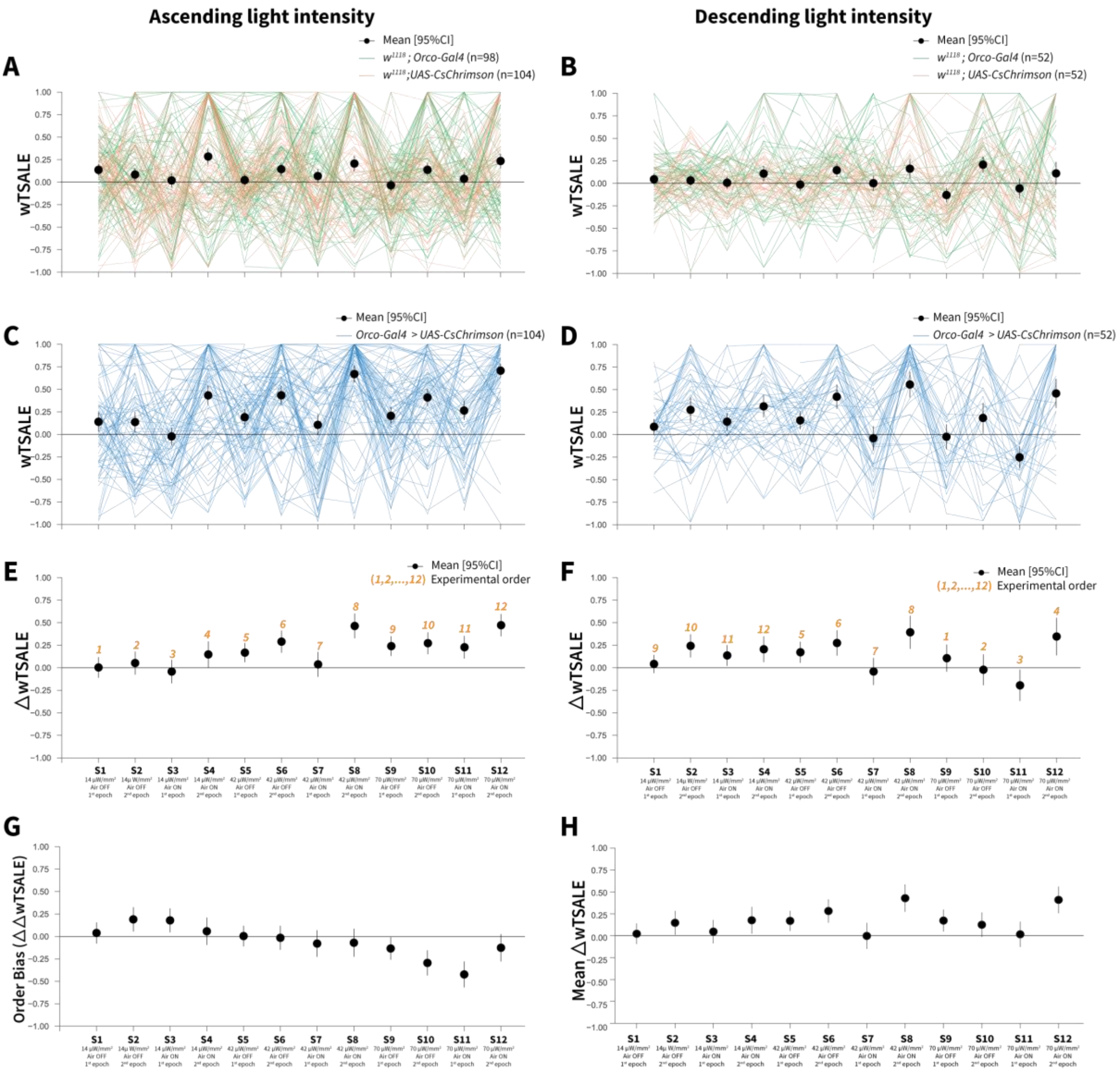


**Figure 2—figure supplement 1. Orco-neuron activation triggers attraction regardless of experimental order**

**A.** The light preference of control flies and the mean wTSALE differences across 12 epochs, in which the light intensities were applied in an ascending order. The green lines represent $w^{1118}$; *Orco-Gal4* flies, and the orange lines represent $w^{1118}$; *UAS-CsChrimson* flies. The black dots represent the mean wTSALE scores (with 95% CIs) per epoch.

**B.** The light preference of the control flies in which the light intensities were applied in a descending order. The green lines represent $w^{1118}$; *Orco-Gal4* flies, and the orange lines represent $w^{1118}$; *UAS-CsChrimson* flies. The black dots represent the mean wTSALE scores (with 95% CIs) per epoch.

**C.** The light preference of the test flies in which the light intensities were applied in an ascending order. The blue lines represent *Orco-Gal4* > *UAS-CsChrimson* flies.

**D.** The test flies' light preference was presented across the 12 epochs. The blue lines represent *Orco-Gal4* > *UAS-CsChrimson* flies.

**E.** The mean wTSALE differences between the test and combined-control flies across the 12 conditions, in ascending order. The orange numbers indicate the order of the respective experiment. The flies were attracted to activity in the *Orco+* neurons in the absence of wind.
**F.** The mean wTSALE differences between the test and combined-control flies across 12 conditions in the descending order. The flies were attracted to activity in the *Orco+* neurons in the absence of wind.
**G.** A comparison plot of the effect sizes from experiments in which the light intensities were applied in an ascending or descending order. The black dots represent the difference of the mean ΔwTSALE differences (ΔΔwTSALE) from the two differently ordered experiments.
**H.** The mean effect sizes of the ascending and descending light-intensity experiments across the epochs. The black dots represent the mean of the ΔwTSALE differences (ΔwTSALE) from the two differently ordered experiments along with 95% CIs.

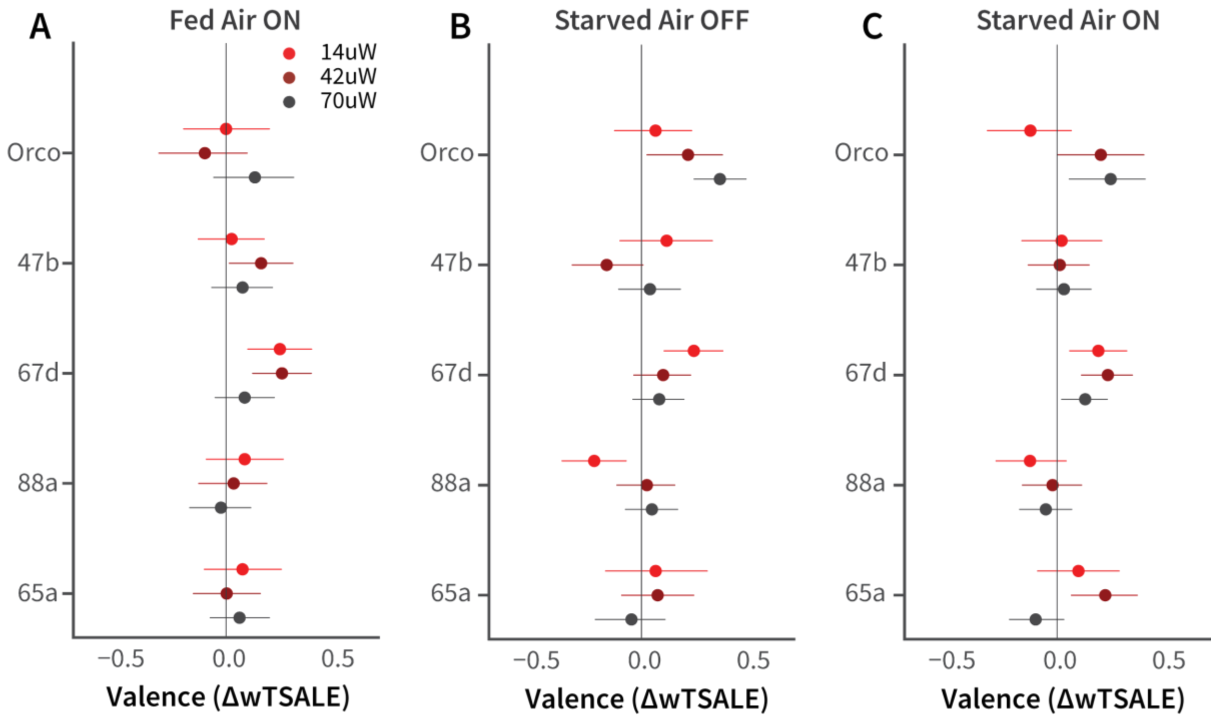


**Figure 2—figure supplement 2. Female-fly responses to activation of Orco and four pheromone-related cells**

**A-C.** The olfactory valence of the *Orco* and four single-ORN types were tested in fed or starved female flies with or without airflow. The valence responses are represented as the mean difference (ΔwTSALE) of control (N ≅ 104) and test (N ≅ 52) flies, along with 95% CIs.

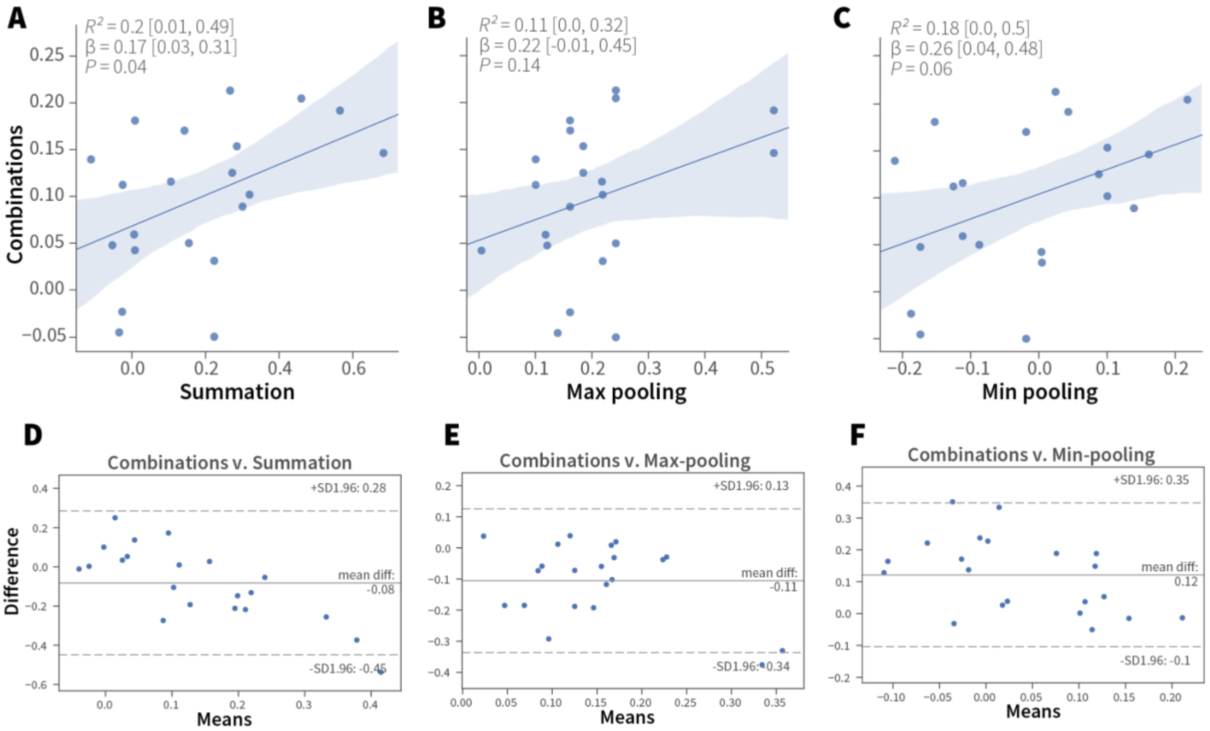


**Figure 5—figure supplement 1. Linear analyses of combination valence results.**

**A-C.** Simple linear regression analyses of the experimental ORN-combo results and the predictions by the (A) summation, $R^2$ = 0.2 [95CI 0.01, 0.49], $P$ = 0.04; (B) max-pooling $R^2$ = 0.11 [95CI 0.00, 0.32], $P$ = 0.14; and (C) min-pooling $R^2$ = 0.18 [95CI 0.0, 0.5], $P$ = 0.06 functions.

**D–F.** Agreement analysis of the observed and predicted ORN-combo valence responses by three pooling functions. Bland-Altman plots of ORN-combo valence responses and predictions by the (D) summation, (E) max-pooling, and (F) min-pooling models. Even though the mean difference of the two tested values (y axis) is low in all models, the limits of agreement (SD –1.96, +1.96) are sufficiently wide for them to be considered dissimilar.

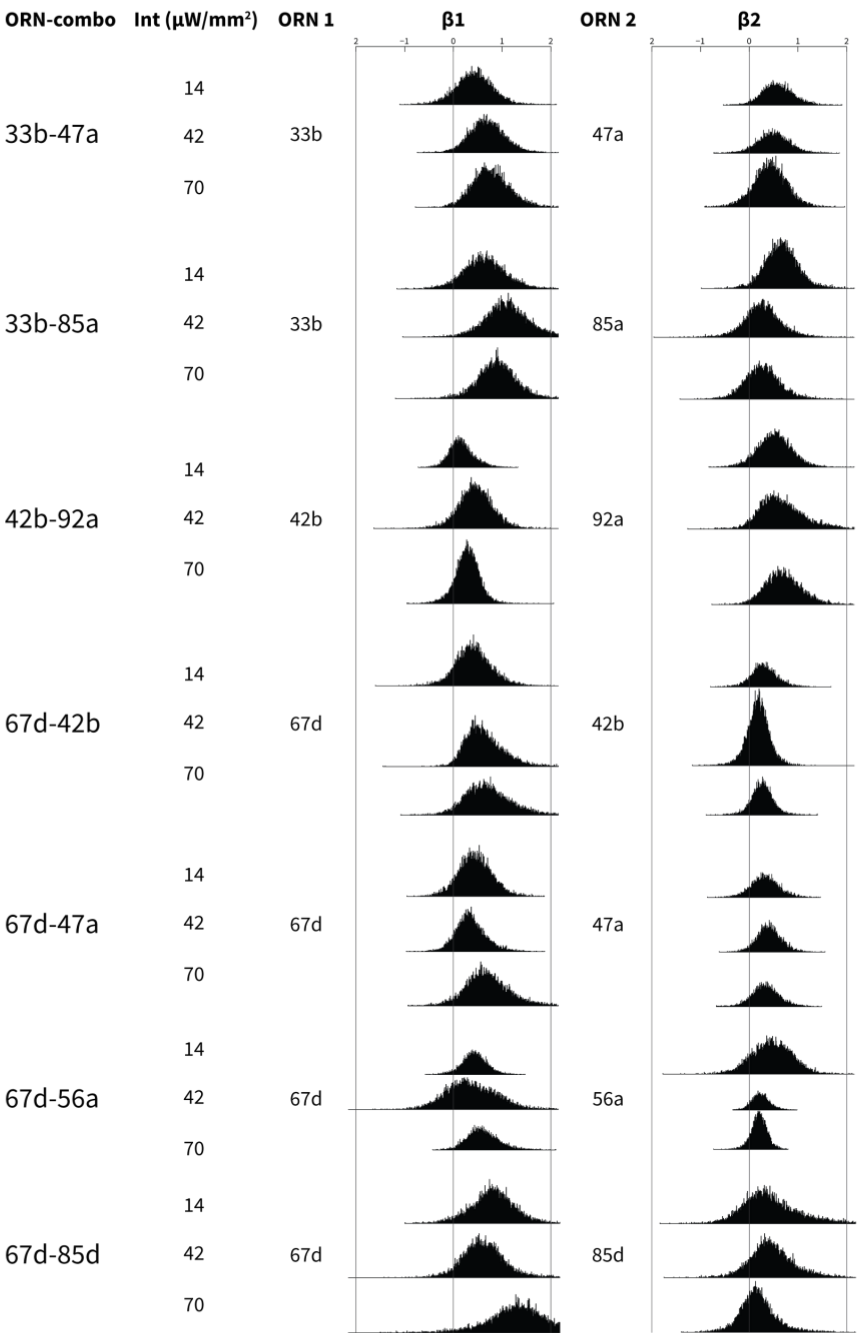


**Figure 5—figure supplement 2. Bootstrapped distributions of β weights of the ORN-combo constituents**

Associations between ORN-combos and their constituent single-ORN types are tested by using a multiple linear regression analysis approach. ORN1 and ORN2 columns indicate the odor receptor types that are used to generate the respective ORN-combo. The light intensity used in the experiments are shown in the intensity (Int; $\mu W/mm^2$) column. β1 and β2 columns present the bootstrapped β distributions of the ORN1 and ORN2, respectively; the y axes indicate the count, and are all plotted using the same scale.

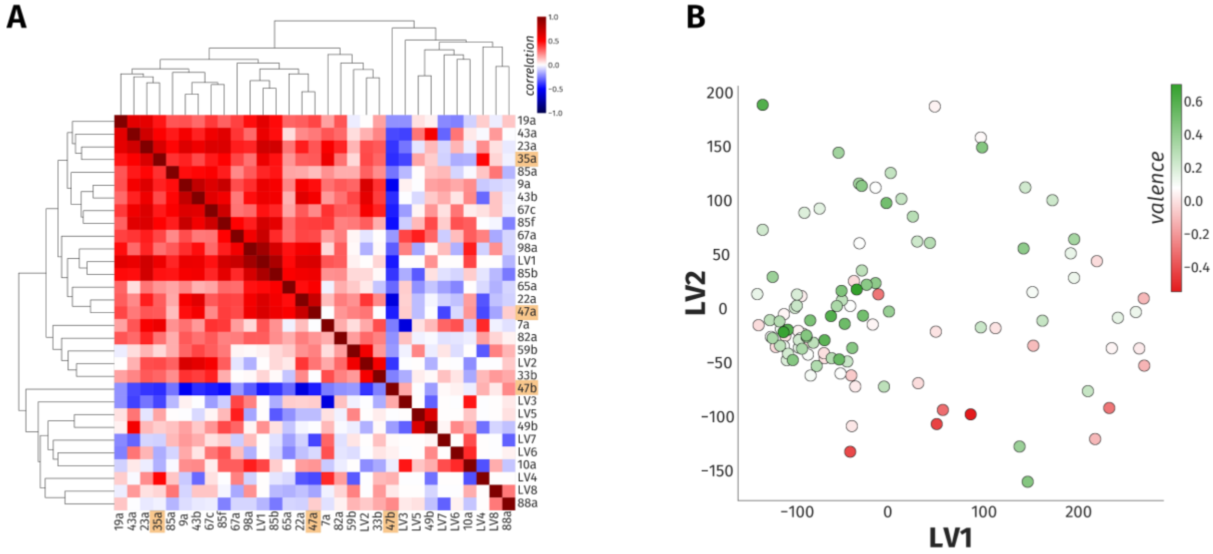


**Figure 5—figure supplement 3. A linear model accounts for only 23% of the variance in odor behavior**

**A.** The heatmap of hierarchical clustering shows the 27 × 27 Pearson correlation coefficients among the 23 ORN types and eight LVs from PLS-DA analysis. The internal correlation of LVs and their constituent ORN-types is indicated by color, where blue and red ends of the spectrum represent negative and positive correlations, respectively. The three ORN types that produced a valence response in the WALISAR screen are highlighted in orange.

**B.** A scatter plot displays projection of 110 odorants to two-dimensional LV space. The valence of the odorants are shown with a color gradient ranging from red to green, indicating aversive and attractive odorants, respectively.

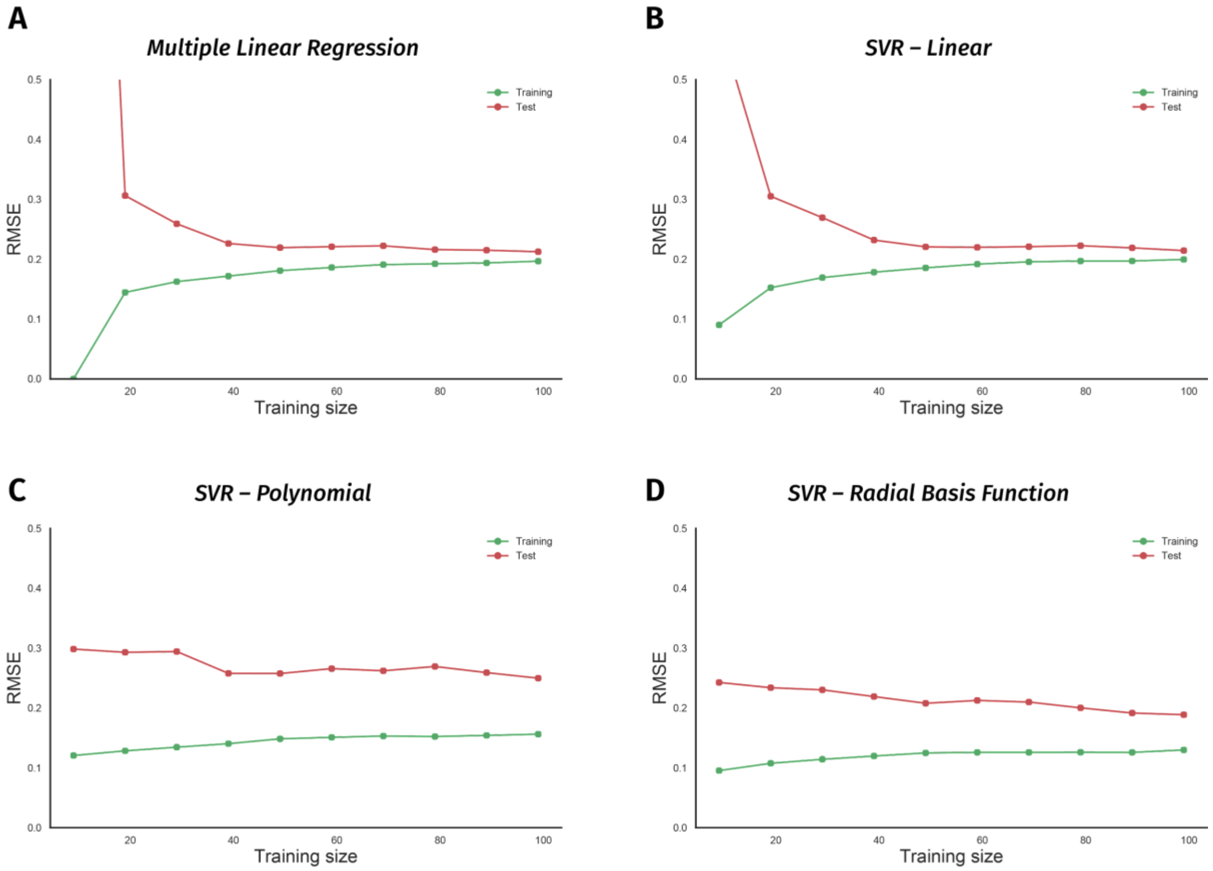


**Figure 5—figure supplement 4. Non-linear models suffer from the small size of the odor-valence data set**

**A.** Performance of a multiple linear regression (MLR) model is shown as the training data size is incrementally increased. The y-axis indicates the root-mean-squared-error (RMSE), while the x-axis is the size of the training data-set. The green and red traces in all the panels represent error rates on training and 10-fold cross-validation test data, respectively.

**B.** Learning curve of a support vector regression (SVR) model with a linear kernel is drawn. Both MLR and SVR linear models show convergent learning curves.

**C.** Performance of a support vector regression (SVR) model using a non-linear, polynomial kernel is plotted over an increasing training data-set size.

**D.** Error rates of a support vector regression (SVR) model with a non-linear, radial-basis-function kernel are plotted over a growing training data size. Both nonlinear models' learning curves fail to converge, indicating overfit.

**Supplementary File 1. ORN types with known innate valence.**
The innate responses of 24 types of ORNs that have been reported at time of writing (January, 2020). The valence response, experimental assay, stimulus type, sex, and the developmental stage of the flies varied across studies. The valence column shows the response direction produced by the respective ORN: negative (-), positive (+), or indifferent (o). The assay column presents the nature of the assay used in the experiments: oviposition (place preference for egg-laying in female flies), two-choice (any assay by which the flies are presented with a choice to activate the ORN), locomotor (the assay in which motor behavior of the larvae is used to deduce valence). The stimulus column shows the type of the stimulus applied on the ORN: olfactogenetics (geosmin on ORNs that ectopically express the Or56a receptor), optogenetic (light stimulus on genetically modified ORNs), odor. In the sex column, F, M, M/F, and N/A indicate female, male, both male and female, and information that is not available, respectively. The stage column presents the developmental stage of the animals used in the experiment: larva, adult, and both (larva and adult).

| **Receptor** | **Valence** | **Assay** | **Stimulus** | **Sex** | **Stage** | **Reference** |
|---|---|---|---|---|---|---|
| Or7a | − | oviposition | olfactogenetics | F | adult | (Chin et al., 2018) |
| Or7a | o | two-choice | optogenetic | M | adult | ***this study*** |
| Or19a | + | oviposition | odor | F | adult | (Dweck et al., 2013) |
| Or19a | o | oviposition | olfactogenetics | F | adult | (Chin et al., 2018) |
| Or19a | o | two-choice | optogenetic | M | adult | ***this study*** |
| Or22a | + | two-choice | odor | M/F | adult | (Knaden et al., 2012) |
| Or22a | + | two-choice | odor | F | adult | (Semmelhack and Wang, 2009) |
| Or22a | − | two-choice | odor | F | adult | (Gao et al., 2015) |
| Or22a | + | two-choice | optogenetic | M | adult | (Bell and Wilson, 2016) |
| Or22a | o | oviposition | olfactogenetics | F | adult | (Chin et al., 2018) |
| Or22a | o | two-choice | optogenetic | M | adult | ***this study*** |
| Or23a | o | oviposition | olfactogenetics | F | adult | (Chin et al., 2018) |
| Or23a | o | two-choice | optogenetic | M | adult | ***this study*** |
| Or35a | o | oviposition | olfactogenetics | F | adult | (Chin et al., 2018) |
| Or35a | + | two-choice | optogenetic | M | adult | ***this study*** |
| Or42a | + | locomotor | optogenetic | NA | larva | (Hernandez-Nunez et al., 2015) |
| Or42a | + | two-choice | odor | NA | larva | (Mathew et al., 2013) |
| Or42a | o | two-choice | odor | F | adult | (Jung et al., 2015) |
| Or42a | + | two-choice | optogenetic | M | adult | (Bell and Wilson, 2016) |
| Or42a | o | oviposition | olfactogenetics | F | adult | (Chin et al., 2018) |

| | | | | | | |
|---|---|---|---|---|---|---|
| Or42a | o | two-choice | olfactogenetics | F | adult | (Chin et al., 2018) |
| Or42a | o | two-choice | optogenetic | M | adult | ***this study*** |
| Or42b | + | two-choice | odor | F | adult | (Semmelhack and Wang, 2009) |
| Or42b | + | two-choice | odor | NA | larva | (Mathew et al., 2013) |
| Or42b | + | two-choice | odor | F | adult | (Gao et al., 2015) |
| Or42b | o | two-choice | odor | F | adult | (Jung et al., 2015) |
| Or42b | + | two-choice | optogenetic | M | adult | (Bell and Wilson, 2016) |
| Or42b | o | oviposition | olfactogenetics | F | adult | (Chin et al., 2018) |
| Or42b | + | two-choice | optogenetic | M | adult | ***this study*** |
| Or43a | o | oviposition | olfactogenetics | F | adult | (Chin et al., 2018) |
| Or43a | o | two-choice | optogenetic | M | adult | ***this study*** |
| Or47a | o | oviposition | olfactogenetics | F | adult | (Chin et al., 2018) |
| Or47a | + | two-choice | optogenetic | M | adult | ***this study*** |
| Or47b | − | oviposition | olfactogenetics | F | adult | (Chin et al., 2018) |
| Or47b | + | two-choice | optogenetic | M | adult | ***this study*** |
| Or49a | − | oviposition | olfactogenetics | F | adult | (Chin et al., 2018)c |
| Or49a | o | two-choice | optogenetic | M | adult | ***this study*** |
| Or56a | − | two-choice | odor | NA | adult | (Stensmyr et al., 2012) |
| Or56a | + | two-choice | optogenetic | M | adult | (Bell and Wilson, 2016) |
| Or56a | − | two-choice | optogenetic | M | adult | ***this study*** |
| Or59c | − | oviposition | olfactogenetics | F | adult | (Chin et al., 2018) |
| Or59c | − | two-choice | optogenetic | M | adult | ***this study*** |
| Or65a | o | oviposition | olfactogenetics | F | adult | (Chin et al., 2018) |
| Or65a | o | two-choice | optogenetic | M | adult | ***this study*** |
| Or67b | + | two-choice | odor | NA | larva | (Mathew et al., 2013) |
| Or67b | + | two-choice | optogenetic | M | adult | (Bell and Wilson, 2016) |
| Or67b | − | oviposition | olfactogenetics | F | adult | (Chin et al., 2018) |
| Or67b | o | two-choice | optogenetic | M | adult | ***this study*** |
| Or67d | o | oviposition | olfactogenetics | F | adult | (Chin et al., 2018) |
| Or67d | + | two-choice | optogenetic | M | adult | ***this study*** |
| Or71a | − | oviposition | olfactogenetics | F | adult | (Chin et al., 2018) |
| Or71a | o | two-choice | olfactogenetics | F | adult | (Chin et al., 2018) |
| Or71a | o | two-choice | optogenetic | M | adult | ***this study*** |
| Or82a | − | oviposition | olfactogenetics | F | adult | (Chin et al., 2018) |
| Or82a | o | two-choice | optogenetic | M | adult | ***this study*** |
| Or83c | + | two-choice | odor | M/F | adult | (Ronderos et al., 2014) |
| Or83c | − | oviposition | olfactogenetics | F | adult | (Chin et al., 2018) |

| | | | | | | |
|---|---|---|---|---|---|---|
| Or83c | + | two-choice | optogenetic | M | adult | ***this study*** |
| Or85a | + | two-choice | odor | M/F | adult | (Knaden et al., 2012) |
| Or85a | − | two-choice | odor | F | adult | (Semmelhack and Wang, 2009) |
| Or85a | − | two-choice | odor | F | adult | (Gao et al., 2015) |
| Or85a | + | two-choice | optogenetic | M | adult | (Bell and Wilson, 2016) |
| Or85a | − | oviposition | olfactogenetics | F | adult | (Chin et al., 2018) |
| Or85a | o | two-choice | olfactogenetics | F | adult | (Chin et al., 2018) |
| Or85a | o | two-choice | optogenetic | M | adult | ***this study*** |
| Or85d | − | oviposition | olfactogenetics | F | adult | (Chin et al., 2018) |
| Or85d | − | two-choice | optogenetic | M | adult | ***this study*** |
| Or88a | o | oviposition | olfactogenetics | F | adult | (Chin et al., 2018) |
| Or88a | o | two-choice | optogenetic | M | adult | ***this study*** |
| Or92a | + | two-choice | odor | F | adult | (Semmelhack and Wang, 2009) |
| Or92a | + | two-choice | optogenetic | M | adult | (Bell and Wilson, 2016) |
| Or92a | o | oviposition | olfactogenetics | F | adult | (Chin et al., 2018) |
| Or92a | o | two-choice | optogenetic | M | adult | ***this study*** |
| Gr21a/Gr63a | − | two-choice | odor | NA | adult | (Suh et al., 2004) |
| Gr21a/Gr63a | − | two-choice | odor | M/F | both | (Faucher et al., 2006) |
| Gr21a/Gr63a | − | two-choice | optogenetic | NA | adult | (Suh et al., 2007) |
| Gr21a/Gr63a | − | two-choice | odor | F | adult | (Poon et al., 2010) |
| Gr21a/Gr63a | − | two-choice | optogenetic | M | adult | (Bell and Wilson, 2016) |
| Gr21a/Gr63a | o | oviposition | olfactogenetics | F | adult | (Chin et al., 2018) |
| Gr21a/Gr63a | o | two-choice | olfactogenetics | F | adult | (Chin et al., 2018) |
| Gr21a/Gr63a | − | two-choice | optogenetic | M | adult | ***this study*** |
| Orco | o | two-choice | optogenetic | NA | adult | *(Suh et al., 2007)* |
| Orco | + | two-choice | optogenetic | M | adult | (Bell and Wilson, 2016) |
| Orco | + | two-choice | optogenetic | M | adult | ***this study*** |

**Supplementary File 2 - Sheet 1. Ten-fold cross-validation of ORN–preference prediction models.**

RMSE scores were calculated with cross-validation for four models of ORN activity and odor preference. All RMSE scores are close to the standard deviation of the dependent variable (**σ** = 0.23).

| Model | RMSE |
|---|---|
| **Multiple Linear Regression** | 0.22 |
| **SVM - Linear** | 0.21 |
| **SVM - Polynomial** | 0.20 |
| **SVM - RBF** | 0.19 |

**Supplementary File 2 - Sheet 2. Ten-fold cross-validation of single-ORN-valence-weighted prediction models.**

| Model | RMSE |
|---|---|
| **Multiple Linear Regression** | 0.22 |
| **SVM - Linear** | 0.22 |
| **SVM - Polynomial** | 0.20 |
| **SVM - RBF** | 0.20 |

**Supplementary File 2 - Sheet 3. A comparison of the experimental designs of two optogenetic studies investigating ORN-valence behaviour.**

| | Bell & Wilson, 2016 | This study |
|---|---|---|
| **Number of ORNs tested** | 8 | 45 |
| **Number of ORN pairs analyzed** | 36 | 7 |
| **Total study N of Gal4 controls** | 0 | ~5148 |
| **Total study N of UAS controls** | 88 | ~5148 |
| **Total study N of test flies** | ~2512 | ~5148 |
| **Independent genetic controls in experiments** | No | Yes |
| **Uses valence effect size** | No | Yes |
| **Number of light intensities** | 8 | 3 |
| **Receptor types assessed for airflow effect** | 1 | 45 |
| **Technical replicates per experiment** | 16 | 1 |
| **Optogenetic light duration (min)** | 64 | 9 |
| **Opsin type** | Channelrhodopsin-2 | CsChrimson |
| **Optogenetic light color** | Blue | Red |
| **Blind flies** | Yes | No |
| **Heat compensation** | Yes | No |
| **Highest optogenetic light intensity (μW/mm$^2$)** | 1500 | 72 |
| **Optogenetic light-induced heat (°C)** | Not reported | 0.3 |
| **Chamber dimensions** | 50 × 5 × 1.2 mm | 50 × 4 × 3 mm |
| **Physiological characterization of stimulation** | Yes | No |
| **Fly transfer to chambers** | Aspiration | Ice anesthesia |

**Supplementary File 2 - Sheet 4. A list of ORN-Gal4 lines used in the study.**

| Gal4 line | Genotype | BDSC stock # |
|---|---|---|
| **Or7a** | *w[*]; P{w[+mC]=Or7a-GAL4.C}214t1.1* | 23908 |
| **Or9a** | *w[*]; P{w[+mC]=Or9a-GAL4.C}106t6.1/TM3, Sb[1]* | 23918 |
| **Or10a** | *w[*]; P{w[+mC]=Or10a-GAL4.F}34.2A; TM2/TM6B, Tb[1]* | 9944 |
| **Or13a** | *w[*]; P{w[+mC]=Or13a-GAL4.C}229t56.2/TM3, Sb[1]* | 23886 |
| **Or19a** | *w[*]; P{w[+mC]=Or19a-GAL4.F}61.2* | 9948 |
| **Or19b** | *w[*]; P{w[+mC]=Or19b-GAL4.C}218t6.1* | 23889 |
| **Or22a** | *w[*]; P{w[+mC]=Or22a-GAL4.7.717}14.2* | 9951 |
| **Or22b** | *w[1118]; P{w[+mC]=Or22b-GAL4.10287}105.2* | 23289 |
| **Or23a** | *w[*]; P{w[+mC]=Or23a-GAL4.7.818}17.2* | 9955 |
| **Or33a** | *w[*]; Bl[1]/CyO; P{w[+mC]=Or33a-GAL4.F}126.4A* | 9962 |
| **Or33b** | *w[*]; Bl[1]/CyO; P{w[+mC]=Or33b-GAL4.F}83.14/TM6B, Tb[1]* | 9963 |
| **Or33c** | *w[*]; P{w[+mC]=Or33c-GAL4.F}78.3* | 9966 |
| **Or35a** | *w[*]; P{w[+mC]=Or35a-GAL4.F}109.3* | 9968 |
| **Or42a** | *w[*]; P{w[+mC]=Or42a-GAL4.F}48.1* | 9969 |
| **Or42b** | *w[*]; P{w[+mC]=Or42b-GAL4.F}64.1* | 9972 |
| **Or43a** | *w[*]; P{w[+mC]=Or43a-GAL4.W}27.6* | 9974 |
| **Or43b** | *w[*]; P{w[+mC]=Or43b-GAL4.C}110t6.3* | 23895 |
| **Or46a** | *w[*]; P{w[+mC]=Or46a-GAL4.1.875}9.10A* | 9979 |
| **Or47a** | *w[*]; P{w[+mC]=Or47a-GAL4.8.239}15.4A* | 9982 |
| **Or47b** | *w[*]; P{w[+mC]=Or47b-GAL4.7.467}15.6* | 9984 |
| **Or49a** | *w[*]; P{w[+mC]=Or49a-GAL4.F}47.2A* | 9985 |
| **Or49b** | *w[*]; P{w[+mC]=Or49b-GAL4.F}80.1* | 9986 |
| **Or56a** | *w[*]; P{w[+mC]=Or56a-GAL4.C}113t53.2* | 23896 |
| **Or59b** | *w[*]; P{w[+mC]=Or59b-GAL4.C}114t2.2* | 23897 |

| | | |
|---|---|---|
| **Or59c** | *w[*]; P{w[+mC]=Or59c-GAL4.C}129t1.1* | 23899 |
| **Or65a** | *w[*]; P{w[+mC]=Or65a-GAL4.F}72.5* | 9993 |
| **Or65b** | *w[*]; P{w[+mC]=Or65b-GAL4.C}198t57.1* | 23902 |
| **Or65c** | *w[*]; P{w[+mC]=Or65c-GAL4.C}221t53.1/TM3, Sb[1]* | 23903 |
| **Or67a** | *w[*]; P{w[+mC]=Or67a-GAL4.C}137t3.3* | 23904 |
| **Or67b** | *w[*]; P{w[+mC]=Or67b-GAL4.F}68.3/TM6B, Tb[1]* | 9995 |
| **Or67c** | *w[*]; P{w[+mC]=Or67c-GAL4.C}116t3.2/CyO* | 23905 |
| **Or67d** | *w[*]; P{w[+mC]=Or67d-GAL4.F}57.2* | 9998 |
| **Or69a** | *w[*]; P{w[+mC]=Or69a-GAL4.F}81.4* | 10000 |
| **Or71a** | *w[*]; P{w[+mC]=Or71a-GAL4.F}30.4* | 23122 |
| **Or82a** | *w[*]; P{w[+mC]=Or82a-GAL4.F}135.1; TM2/TM6B, Tb[1]* | 23125 |
| **Or83c** | *w[*]; P{w[+mC]=Or83c-GAL4.F}73.4A* | 23132 |
| **Or85a** | *w[*]; P{w[+mC]=Or85a-GAL4.F}67.4* | 24461 |
| **Or85b** | *w[*]; P{w[+mC]=Or85b-GAL4.C}179t55.1* | 23912 |
| **Or85d** | *w[*]; P{w[+mC]=Or85d-GAL4.C}143t2.1* | 24148 |
| **Or85e** | *w[1118]; P{w[+mC]=Or85e-GAL4.W}2.19.1* | 23293 |
| **Or85f** | *w[*]; P{w[+mC]=Or85f-GAL4.F}44.6* | 23136 |
| **Or88a** | *w[*]; P{w[+mC]=Or88a-GAL4.F}52.1* | 23137 |
| **Or92a** | *w[*]; P{w[+mC]=Or92a-GAL4.F}62.1* | 23139 |
| **Or98a** | *w[*]; P{w[+mC]=Or98a-GAL4.F}115.1* | 23141 |
| **Orco** | *w[*]; P{w[+mC]=Orco-GAL4.W}11.17; TM2/TM6B, Tb[1]* | 26818 |
